# Clarifying intercellular signalling in yeast: *Saccharomyces cerevisiae* does not undergo a quorum sensing-dependent switch to filamentous growth

**DOI:** 10.1101/2021.10.25.462316

**Authors:** Michela Winters, Violetta Aru, Kate Howell, Nils Arneborg

**Affiliations:** School of Agriculture and Food, Faculty of Veterinary and Agricultural Science, University of Melbourne, Parkville 3010, Australia; Department of Food Science, University of Copenhagen, Frederiksberg 1958, Denmark

**Keywords:** *Saccharomyces cerevisiae*, filamentous growth, budding pattern, signalling metabolites, cell density

## Abstract

*Saccharomyces cerevisiae* can alter its morphology to a filamentous form associated with unipolar budding in response to environmental stressors. Induction of filamentous growth is suggested under nitrogen deficiency in response to alcoholic signalling molecules through a quorum sensing mechanism. To investigate this claim, we analysed the budding pattern of *S. cerevisiae* cells over time under low nitrogen while concurrently measuring cell density and extracellular metabolite concentration. We found that the proportion of cells displaying unipolar budding increased between local cell densities of 4.8×10^6^ and 5.3×10^7^ cells/ml. However, the observed increase in unipolar budding could not be reproduced when cells were prepared at the critical cell density and in conditioned media. Removing the nutrient restriction by growth under high nitrogen conditions also resulted in an increase in unipolar budding between local cell densities of 5.2×10^6^ and 8.2×10^7^ cells/ml, but there were differences in metabolite concentration compared to the low nitrogen conditions. This suggests that neither cell density, metabolite concentration, nor nitrogen deficiency were necessary or sufficient to increase the proportion of unipolar budding cells. It is therefore unlikely that quorum sensing is the mechanism controlling the switch to filamentous growth in *S. cerevisiae*. Only a high concentration of the putative signalling molecule, 2-phenylethanol resulted in an increase in unipolar budding, but this concentration was not physiologically relevant. We suggest that the compound 2-phenylethanol acts through a toxicity mechanism, rather than quorum sensing, to induce filamentous growth.

**IMPORTANCE:** Investigating dimorphism in the model organism *Saccharomyces cerevisiae* has been instrumental in understanding the signalling pathways that control hyphal growth and virulence in human pathogenic fungi. Quorum sensing was proposed to signal morphogenesis in *S. cerevisiae* populations. This mechanism requires the switch to filamentous growth to occur at a critical quorum sensing molecule concentration corresponding to a critical cell density. However, evidence for this mechanism is sparse and limited by the use of non-physiologically relevant concentrations of signalling metabolites. Our study designed a methodology to address this gap and may be applied to further studies of dimorphism in other types of yeasts. A significant implication of our findings is that morphogenesis does not occur in *S. cerevisiae* via a quorum sensing mechanism, and this important definition needs to be corrected. Mechanistic studies to understand dimorphism in yeasts, by considering metabolite concentrations, will further shed light onto this important cellular behaviour.

## INTRODUCTION

The yeast *Saccharomyces cerevisiae* has the ability to exist in different multicellular forms depending on its environment. In liquid environments, the yeast cells can be sessile or planktonic and may form flocs and flors, and in solid and semi-solid substrates, colonies, biofilms, filaments and mats are observed (1). Filamentous growth of *S. cerevisiae* commonly occurs under nutrient starvation and has been hypothesised as a mechanism to allow the cells to grow into their immediate environment and forage for available nutrients (2, 3). This morphogenesis is described as pseudohyphal or invasive depending on whether the cells exist as diploids or haploids and can be observed both at a colony and cellular level (2, 4–9). When growing on agar plates, colony morphology and invasion reveal filamentous growth, and these macro-observations are determined by budding pattern and cell shape that indicate filamentous growth at the cellular level (2, 4–6, 10, 11).

Budding is the method by which *S. cerevisiae* cells divide during their mitotic cell cycle. The budding pattern is determined by the orientation of new buds with respect to the mother-daughter junction and can be differentiated as axial, bipolar or unipolar (Fig. 1) (2, 5, 12–15). Whether the cell is a diploid or haploid will determine which budding pattern is exhibited when cells grow planktonically (12, 13). Haploid cells bud in an axial manner whereby the new bud consistently emerges adjacent to or overlapping the pole of the previous mother-daughter junction. Diploid cells, however, bud according to a bipolar pattern, where buds can emerge either adjacent to the mother-daughter junction or at the opposite pole. Usually, the first bud site of a new daughter cell has a strong opposite pole bias, while second and subsequent buds have no bias towards a pole (2, 12, 14, 15). Both haploids and diploids transition to a unipolar budding pattern when cells switch to filamentous growth. This pattern occurs when the daughter buds consistently from the pole opposite the junction to its mother (2, 5, 7–10). Cell division is thus polarized by the serial reiteration of unipolar budding away from the mass of cells in the colony into chains of interconnected cells (2).

**FIG 1.**
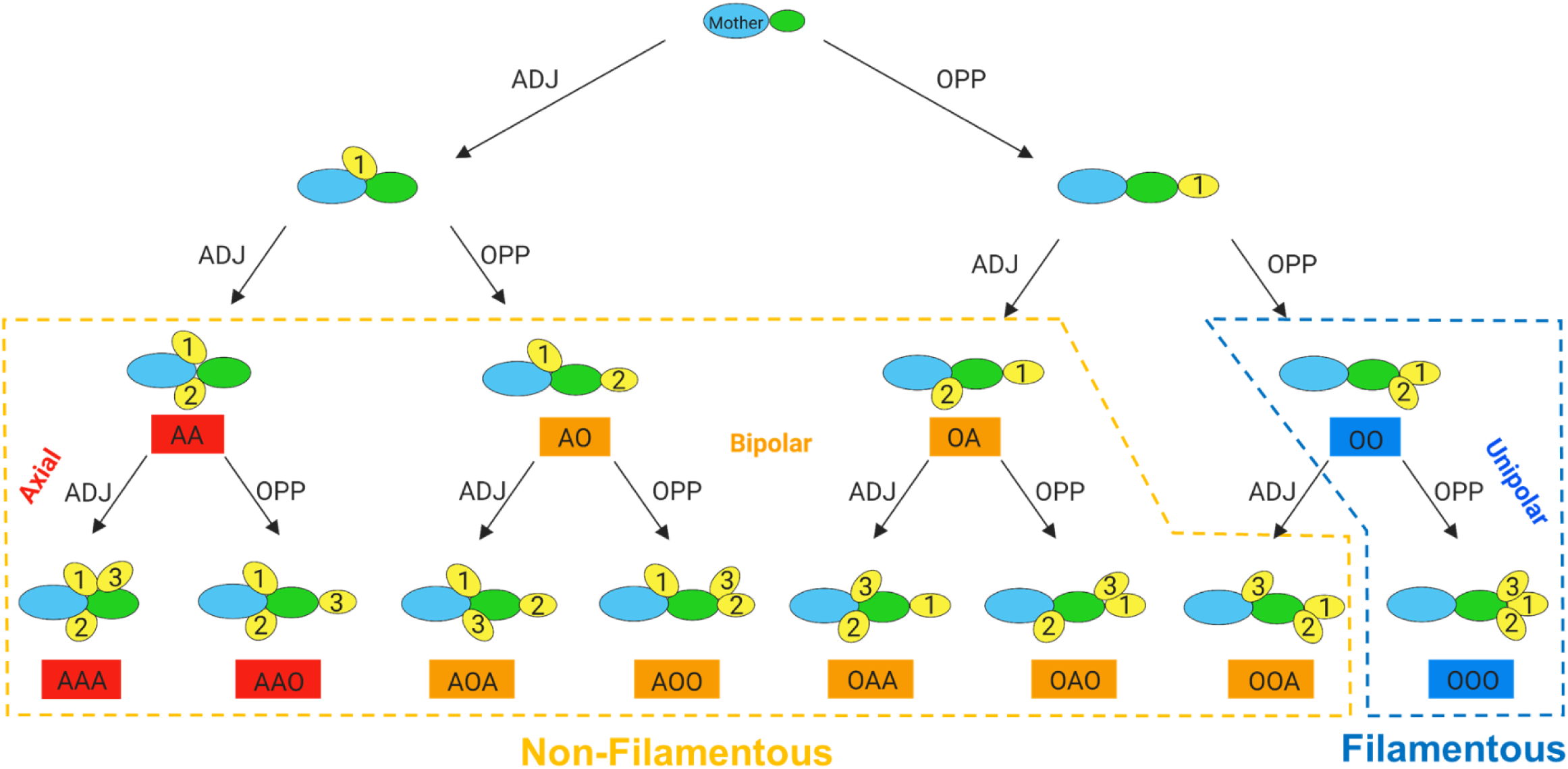
Schematic diagram of how the budding pattern is obtained from the bud site sequence of a new daughter cell. Bud site sequence is assigned for new daughters with both two and three bud sites and categorised to its corresponding budding pattern as shown. For cells growing in non-filamentous form, haploids will adopt an axial pattern (red). This pattern is defined as new buds emerging consistently adjacent to the junction to the mother. Diploids will adopt a bipolar pattern (orange) whereby buds may emerge either adjacent or opposite the junction to the mother. When filamentous growth occurs, both cell types transition to a unipolar budding pattern (blue). This pattern is defined as new buds emerging consistently from the end opposite the junction to the mother. In this way, budding pattern can be used as a proxy for tracking filamentous growth in liquid media (2, 5, 12–15).

*S. cerevisiae* cells undergoing filamentous growth have a higher proportion of unipolar budding compared to non-filamentous cells. Previous research established that when two bud sites are observed, non-filamentous cells show 70-71% unipolar budding, while filamentous cells show 90-100% unipolar budding (2, 5). For cells with three bud sites, this proportion of unipolar budding was observed to be 26% for non-filamentous cells and 97% for filamentous cells (5). Furthermore, under filamentous growth inducing conditions (nitrogen deficiency), unipolar budding was found to increase by an average of 35% (16). Observation of a higher proportion of unipolar budding cells thus signifies that cells are on track to grow as filaments.

Filamentous growth is induced in *S. cerevisiae* cells in response to different environmental cues and stressors, including nutrient deficiency and signalling molecules. Filamentous cells are considered more able to scavenge nutrients when exposed to a variety of extracellular stressors (17). Nutrient starvation, including lack of nitrogen (2, 9, 18–20), can trigger the morphological change to filamentous growth. An additional adaptation to nitrogen starvation is the secretion of autoregulatory aromatic alcohols that stimulate filamentous growth through a quorum sensing mechanism (18, 21–23).

Quorum sensing is a form of intercellular signalling where interactions between organisms are mediated by intercellular signalling molecules (24). Quorum sensing involves a cell-density dependent regulation of a behaviour that occurs after a critical signal concentration of quorum sensing molecules is achieved (25). Quorum sensing molecules are excreted and accumulate in the extracellular environment proportional to cell density. At the critical cell density a corresponding threshold concentration of quorum sensing molecules is reached and a synchronised response is triggered in the community of cells (26–31). A prior study observed a quorum sensing-controlled mechanism for the switch to filamentous growth in *S. cerevisiae,* connecting nutrient sensing with aromatic alcohol secretion. Nitrogen and cell density were demonstrated to regulate the production of three auto-induced aromatic alcohols via a positive feedback loop. These included tyrosol (produced in the range 1-8 μM), 2-phenylethanol (in the range of 1-8 μM) and tryptophol (in the range of 1-2 μM). 2-phenylethanol and tryptophol, but not tyrosol, were then found to stimulate filamentous growth when added exogenously at ≥20 μM under nitrogen limiting conditions (18). Further studies have linked exogenous addition of different alcohols to induction of morphogenesis under both high (32) and low nitrogen (9) conditions. These include 1-butanol (9), isoamylol (9, 32), 1-propanol (9, 32) and n-hexanol (32). However, all these studies induced morphogenesis by using exogenous alcohol concentrations much higher than those found to be produced when the cell is growing in defined or undefined media (25).

Intercellular signalling interactions need to be defined by specific criteria to be accurately identified and understood. Previous definitions of quorum sensing have tended to be very broad so that any morphological behaviours in yeast may have been inaccurately categorised as being a result of quorum sensing. In our previous work, we set out a series of criteria for what constitutes an intercellular signalling molecule and quorum sensing molecule to promote clarity in classifying microbial interactions (25). When these defined criteria were applied to studies of quorum sensing-controlled filamentous growth in *S. cerevisiae* we found that two criteria were consistently not met in prior research (25). Firstly, there is a lack of evidence that the switch in cellular behaviour occurs at a critical quorum sensing molecule concentration corresponding to a critical cell density. Secondly, no studies used physiologically relevant concentrations of intercellular signalling molecules to trigger the switch to filamentous growth.

In the current study, we aimed to address these two missing criteria on the role of quorum sensing in the switch to filamentous growth in *S. cerevisiae*. We designed a new methodology which uses budding pattern as a proxy for filamentous growth. This involved observing an individual cell, growing over time, to determine the orientation of buds from new daughter cells. By using an oCelloScope^TM^ to capture time-lapse images of budding yeast growing in liquid media, we observed a switch to a filamentous budding pattern. Since our method simultaneously quantified the budding pattern, cell density and metabolite concentration under physiological conditions, we successfully identified the critical cell density and metabolite concentration at which cells switched to filamentous growth. An untargeted metabolome approach through proton (^1^H) NMR meant both previously reported and putative quorum sensing molecules could be monitored. Next, we individually assessed the role of cell density, metabolites, and nitrogen concentration in triggering this switch to filamentous growth. Additionally, we investigated the role of exogenous addition of 2-phenylethanol, a putative quorum sensing molecule, in morphogenesis. Together, our findings suggest that *S. cerevisiae* does not undergo a quorum sensing-dependent switch to filamentous growth, and that 2-phenylethanol acts through a toxicity mechanism rather than as a quorum sensing molecule. Finally, the novel methodology developed here opens the way for further studies of dimorphism in other strains and species of yeasts.

## RESULTS

### *S. cerevisiae* Σ1278b displays a switch in budding pattern under low nitrogen

We designed a time series experiment such that *S. cerevisiae* strains could grow in the same media to obtain an accurate link between our three factors of interest: budding pattern, cell density and metabolite concentration. The cropped images obtained from the oCelloScope^TM^ were analyzed manually and the bud site sequence of new daughters were recorded followed by a classification of their corresponding budding pattern (Fig. 1). A representative example of this assignment process can be seen in Figure 2.

**FIG 2.**
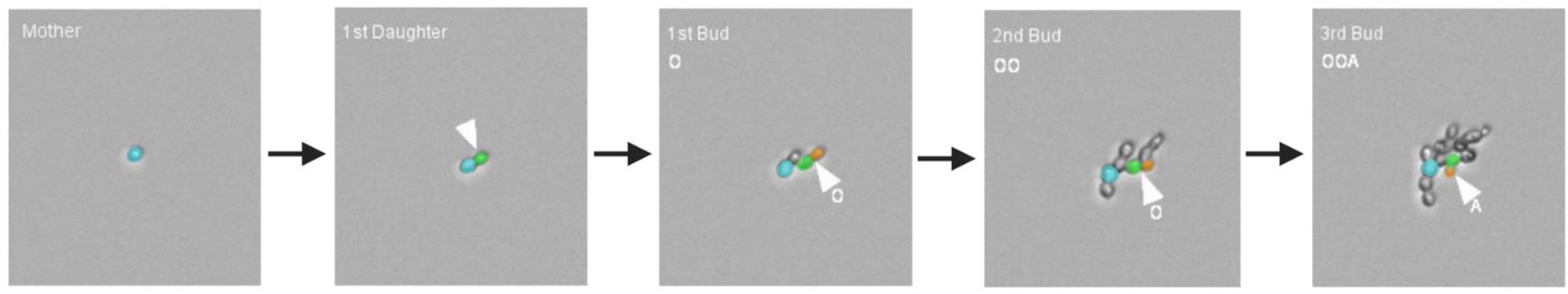
Representative cropped time-lapse images of a new daughter cell being assigned a budding pattern by using its bud site sequence. Time-lapse images were obtained with the oCelloScope^TM^. For the 1st daughter (green), the first bud (orange) emerges opposite the birth end (O), followed by a second bud (orange) also opposite its birth end (OO) and finally the third bud (orange) emerges adjacent to the birth end (OOA). Thus, this daughter would have a bud site sequence of OOA which categorizes it as bipolar budding pattern according to Fig 1.

The growth of *S. cerevisiae* strains S288c and Σ1278b were observed under low nitrogen conditions over 30 hours (Fig. 3). Both S288c and Σ1278b strains grew exponentially up to 20 hours after which their growth slightly declined (Fig. 3A).

**FIG 3.**
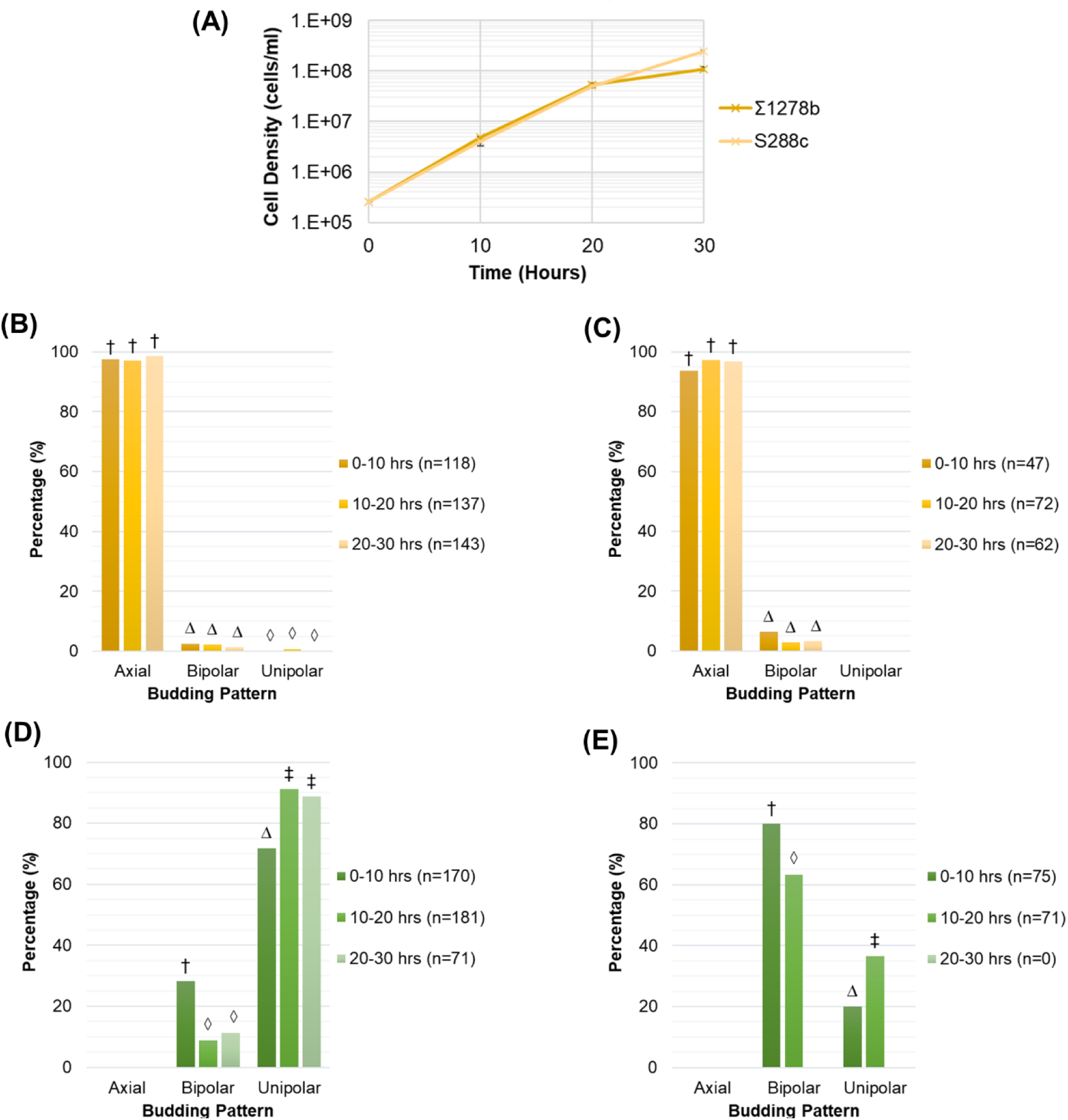
*S. cerevisiae* strain Σ1278b shows an increase in the filamentous unipolar budding pattern under low nitrogen conditions between local cell densities of 4.8×106 and 5.3×107 cells/ml. (A) Change in local cell density under low nitrogen conditions measured by cell counting with a hemocytometer. Means and standard deviation for three biological replicates are shown. (B) Budding pattern for S288cs cells with two bud sites. (C) Budding pattern for S288cs cells with three bud sites. (D) Budding pattern for Σ1278b cells with two bud sites. (E) Budding pattern for Σ1278b cells with three bud sites. The percentage of total cells exhibiting each budding pattern are shown. Budding pattern of cells during growth were determined by manual image analysis of time-lapse images obtained using the oCelloScope^TM^. Different symbols indicate a P value of ≤0.05 as determined by Fishers Exact test.

A predominantly axial budding pattern was observed for the negative control strain S288c for both the two and three bud sites, with no significant changes in budding pattern observed over time (Fig. 3B). Strain Σ1278b budded in a predominantly bipolar or unipolar pattern. Additionally, the proportion of unipolar budding was observed to increase over time, with a corresponding decrease in bipolar budding, for both two and three bud sites (Fig. 3D and E). An increase in the proportion of unipolar budding from 72% to 91% (Fig. 3D) was observed in cells with two bud sites. For cells with three bud sites, it increased from 20% to 36% (Fig. 3E). Due to the growth limitation, most probably caused by nitrogen starvation between 20 and 30 hours (Table S1), the cells were only able to achieve two bud sites and therefore budding pattern data for three bud sites were not available during this period (Fig. 3E). The significant (P≤ 0.05) increase in filamentous unipolar budding occurred between the local cell densities of 4.8×10^6^ and 5.3×10^7^ cells/ml within 10 and 20 hours of growth. These values signify the critical cell density for the switch to filamentous growth.

### Physiological concentrations of metabolites present at the critical cell density and change in budding pattern

To correlate the change in concentration of metabolites over time to the changes in budding pattern and cell density observed, media from the time series experiments were collected, measured by ^1^H NMR spectroscopy, and compounds identified and quantified with SigMa (33). In total, 30 compounds were identified in the media with concentrations quantified for each time period (Table 1). Tryptophol was not identified in our samples though we specially looked for it during the NMR identification stage. 2-phenylethanol and tyrosol were present but did not show a significant increase in concentration until 30 hours with very low concentrations prior to that (Table 1).

**TABLE 1:** Heatmap of the physiological concentrations of compounds detected under low nitrogen conditions during growth of *S. cerevisiae* strain Σ1278b.^a^

| Low Concentration Metabolites (μM) | | | | |
| --- | --- | --- | --- | --- |
| Metabolite Name | 0 hrs | 10 hrs | 20 hrs | 30 hrs |
| 1-Propanol | 0.8 ± 0.1 | 1.4 ± 0.3 | 3.7 ± 0.3 | 6.7 ± 0.5 |
| 2-Hydroxyisovalerate | 0.6 ± 0.2 | 0.7 ± 0.2 | 1.3 ± 0.8 | 2.9 ± 0.2 |
| 2-Oxoglutarate | 3.4 ± 1.2 | 4.4 ± 0.2 | 9.3 ± 2.5 | 61.7 ± 1.3 |
| 2-Phenylethanol | 0.5 ± 0.1 | 0.7 ± 0.2 | 1.5 ± 0.8 | 4.3 ± 0.4 |
| 3-Methylxanthine | 5.9 ± 0.8 | 3.9 ± 0.1 | 4.2 ± 0.4 | 3.7 ± 0.2 |
| Acetaldehyde | 0.4 ± 0.1 | 2.3 ± 0.3 | 19.2 ± 0.6 | 30.0 ± 3.8 |
| Acetoin | 0.4 ± 0.1 | 6.7 ± 4.7 | 8.0 ± 2.1 | 20.1 ± 1.3 |
| Choline | 2.3 ± 0.2 | 2.9 ± 0.9 | 2.0 ± 0.8 | 1.6 ± 0.4 |
| Diacetyl | 5.0 ± 0.4 | 5.4 ± 0.4 | 4.8 ± 0.8 | 5.0 ± 0.1 |
| Fumarate | 0.6 ± 0.1 | 0.5 ± 0.2 | 1.1 ± 0.5 | 14.6 ± 1.4 |
| Hydroxyacetone | 3.6 ± 1.6 | 4.2 ± 1.7 | 2.4 ± 0.2 | 3.3 ± 0.1 |
| Isoamylol | 0.6 ± 0.2 | 0.6 ± 0.4 | 1.0 ± 0.7 | 11.1 ± 1.1 |
| Isopropanol | 0.3 ± 0.1 | 0.4 ± 0.1 | 2.1 ± 0.2 | 4.8 ± 0.7 |
| Malate | 2.4 ± 0.6 | 2.9 ± 0.4 | 6.8 ± 3.7 | 10.2 ± 0.7 |
| Methionine | 0.2 ± 0.1 | 0.2 ± 0.1 | 0.6 ± 0.1 | 1.0 ± 0.1 |
| Myoinositol | 47.3 ± 1.0 | 43.6 ± 7.3 | 33.1 ± 13.0 | 41.4 ± 0.7 |
| Niacin | 9.5 ± 0.2 | 7.8 ± 1.6 | 7.6 ± 2.9 | 9.6 ± 0.2 |
| Pantothenate | 4.3 ± 0.2 | 4.8 ± 0.4 | 5.3 ± 0.6 | 6.0 ± 0.1 |
| Pyridoxine | 7.8 ± 0.2 | 8.3 ± 0.3 | 7.8 ± 0.3 | 7.4 ± 0.1 |
| Pyruvate | 4.9 ± 0.9 | 5.0 ± 0.1 | 30.6 ± 1.5 | 52.8 ± 1.9 |
| Succinate | 1.8 ± 0.2 | 2.5 ± 0.1 | 12.0 ± 7.3 | 45.0 ± 0.2 |
| Trehalose | 91.3 ± 0.5 | 88.1 ± 8.3 | 66.9 ± 26.6 | 85.9 ± 1.3 |
| Tyrosol | 0.2 ± 0.1 | 0.3 ± 0.1 | 0.8 ± 0.2 | 4.9 ± 0.2 |
| Xanthine | 0.9 ± 0.1 | 1.3 ± 0.3 | 0.6 ± 0.1 | 6.6 ± 0.1 |

0 μM 92 μM
| High Concentration Metabolites (μM) | | | | |
| --- | --- | --- | --- | --- |
| Metabolite Name | 0 hrs | 10 hrs | 20 hrs | 30 hrs |
| Acetate | 9 ± 1 | 17 ± 1 | 160 ± 6 | 411 ± 12 |
| Ethanol | 189 ± 40 | 369 ± 8 | 5403 ± 160 | 14334 ± 1023 |
| Formate | 221 ± 2 | 238 ± 2 | 231 ± 9 | 193 ± 4 |
| Glucose ($\times 10^3$) | 107 ± 1 | 103 ± 1 | 115 ± 5 | 99 ± 1 |
| Lactate | 8 ± 2 | 29 ± 1 | 224 ± 8 | 343 ± 38 |

0 μM
115 mM
<sup>a</sup>Metabolites identified through <sup>1</sup>H-NMR and quantified using the SigMa programme (Khakimov et al. 2020). Values are expressed as mean ± SE for two or three biological replicates.

Metabolites potentially involved in the budding pattern switch were chosen from the identified compound list as those that were not present as nutrients in the raw media and had concentrations that increased during growth. The metabolite concentrations that corresponded to the observed switch in budding pattern were 1-propanol (1.4 – 3.7 µM), 2-hydroxyisovalerate (0.7 – 1.3 µM), 2-oxoglutarate (4.4 – 9.3 µM), 2-phenylethanol (0.7 – 1.5 µM), acetaldehyde (2.3 – 19.2 µM), acetate (17 - 160 µM), acetoin (6.7 – 8.0 µM), ethanol (369 – 5403 µM), fumarate (0.5 – 1.1 µM), isoamylol (0.6 – 1.0 µM), isopropanol (0.4 – 2.1 µM), lactate (29 – 224 µM), malate (2.9 – 6.8 µM), pyruvate (5.0 – 30.6 µM), succinate (2.5 – 12.0 µM), and tyrosol (0.3 – 0.8 µM). These values represent the physiological metabolite concentration present at the critical cell density at which cells switch to filamentous growth.

### Critical cell density does not recapitulate the switch in budding pattern

To establish whether the increase in unipolar budding was dependent on cell density alone, we inoculated cells at the critical cell density, recorded at 10 hours, into low nitrogen media (Fig. 4). This cell density was chosen as it was at the beginning of the time period where we observed the significant increase in unipolar budding. At the critical cell density, 71% of cells with two bud sites had a unipolar pattern (Fig. 4A). This was analogous to 72% unipolar budding observed between 0 to 10 hours in the time series experiment (Fig. 3D). This parallel was also seen for cells with three bud sites at the critical cell density where we measured 23% unipolar budding (Fig. 4B) compared to 20% between 0 to 10 hours in the time series results (Fig. 3E). Since the critical cell density did not replicate the increase in unipolar budding, these findings demonstrate that cell density alone is not responsible for the behaviour.

**FIG 4.**
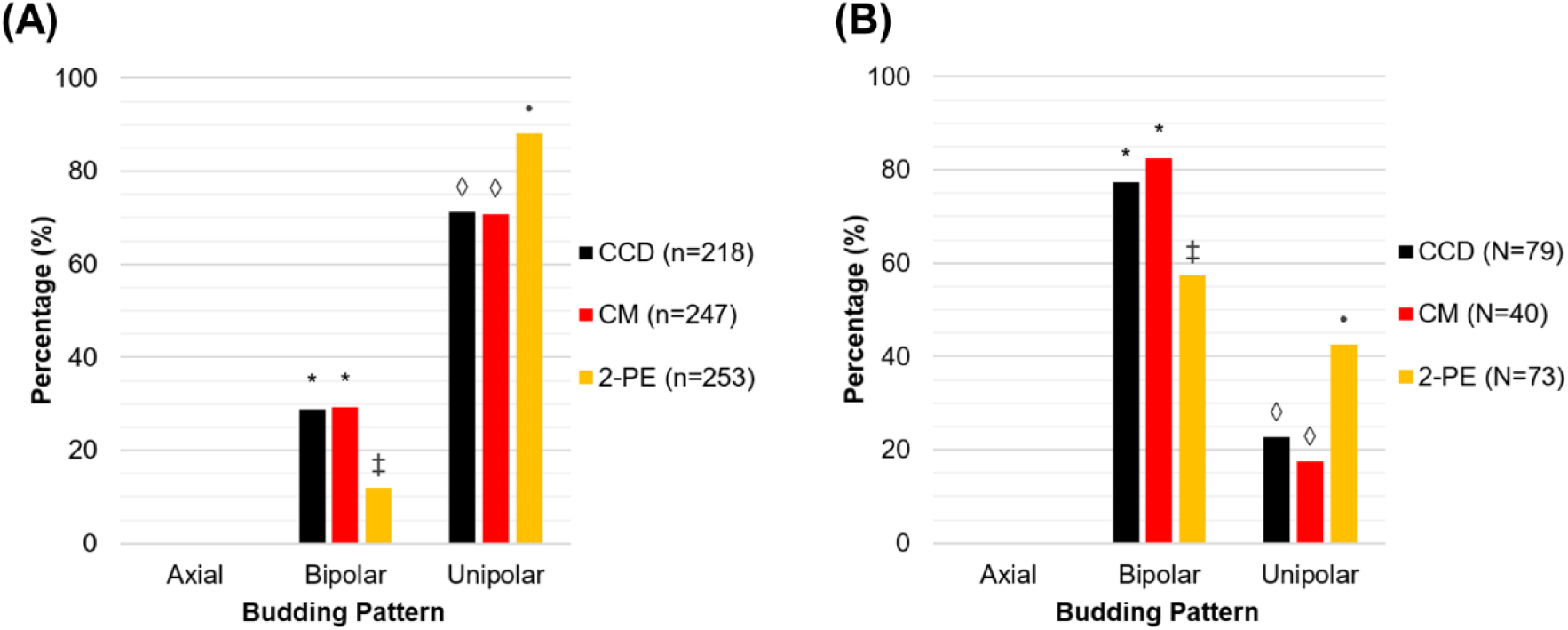
High-concentration 2-phenylethanol recapitulates the increase in unipolar budding, while cells at the critical cell density and growing in conditioned media do not. Experiments were carried out using *S. cerevisiae* strain Σ1278b under low nitrogen condition at the critical cell density to investigate the effect of cell density alone (CCD), the combination of cell density and signalling metabolites (CM), and non-physiological 2-phenylethanol (2PE) on the budding pattern. (A) Budding pattern for Σ1278b cells with two bud sites. (B) Budding pattern for Σ1278b cells with three bud sites. The percentage of total cells exhibiting each budding pattern are shown. Different symbols indicate a P value of ≤0.05 as determined by Fishers Exact test.

### Conditioned media and cells at the critical cell density do not recapitulate the switch in budding pattern

To test whether the combination of physiological metabolites and the critical cell density was a trigger for the increase in unipolar budding, we determined the budding pattern of cells growing in conditioned media from 20 hours of growth at the critical cell density in low nitrogen media (Fig. 4). Growth of cells with two bud sites in conditioned media had 71% unipolar budding (Fig. 4A) and cells with three bud sites had 18% (Fig. 4B). The proportion of unipolar budding observed between the critical cell density and conditioned media conditions were not significantly different; both had 71% for two bud sites (Fig. 4A), and 20% and 18% respectively for three bud sites (Fig. 4B). In both cases the values paralleled the proportion of unipolar budding observed between 0 to 10 hours in the time series experiment; 72% for two bud sites (Fig. 3D) and 20% for three bud sites (Fig. 3E). Thus, the combination of critical cell density and physiological metabolite concentrations were also not accountable for the observed increase in unipolar budding.

### Growth under high nitrogen conditions replicates the switch budding pattern but with different produced metabolite concentrations

To determine whether budding pattern behaviour over time was impacted by nitrogen conditions, we carried out the time series experiment in high nitrogen media. Evidence for the differences in the nitrogen concentration between the two conditions is provided in Table S1.

Similar to growth in low nitrogen, S288c and Σ1278b cells grew exponentially from 0 to 20 hours of the experiment, followed by a slight decrease between 20 and 30 hours (Fig. 5A). However, there was a less pronounced decrease in growth rate for the Σ1278b compared to the S288c under high nitrogen conditions compared to low nitrogen conditions.

**FIG 5.**
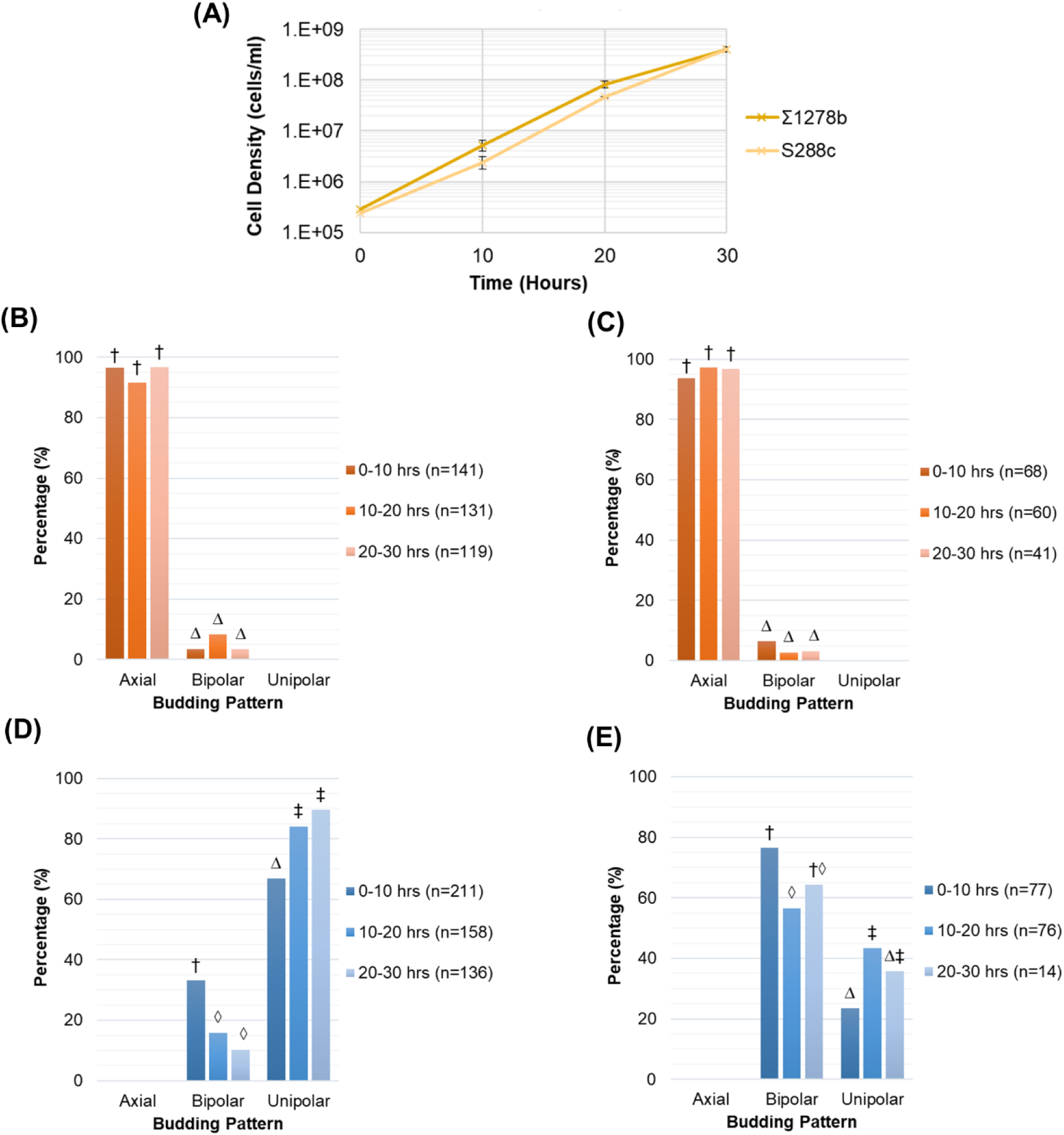
*S. cerevisiae* strains growing under high nitrogen conditions display the same budding pattern behaviour as low nitrogen conditions, with an increase in unipolar budding occurring between local cell densities of 5.2×10^6^ and 8.2×10^7^ cells/ml. (A) Change in local cell density under high nitrogen conditions measured by cell counting with a hemocytometer. Means and standard deviation for three biological replicates are shown. (B) Budding pattern for S288cs cells with two bud sites. (C) Budding pattern for S288cs cells with three bud sites. (D) Budding pattern for Σ1278b cells with two bud sites. (E) Budding pattern for Σ1278b cells with three bud sites. The percentage of total cells exhibiting each budding pattern are shown. Budding pattern of cells during growth were determined by manual image analysis of time-lapse images obtained using the oCelloScope^TM^. Different symbols indicate a P value of ≤0.05 as determined by Fishers Exact test.

Budding pattern behaviour was not significantly different between the two nitrogen conditions. Just as in low nitrogen, S288c showed a predominantly axial budding pattern with no changes over time (Fig. 5B and C) and Σ1278b showed a consistently bipolar or unipolar budding pattern (Fig. 5D and E). Again, we saw a significant increase to a unipolar budding pattern accompanied by a significant decrease in bipolar budding between 10 and 20 hours for cells with both two and three bud sites (Fig. 5D and E). For cells with two bud sites we saw an increase in unipolar budding from 67% to 84% (Fig. 5D). For cells with three bud sites, it increased from 23% to 43% (Fig. 5E). The increase in unipolar budding was associated with local cell densities of 5.2×10^6^ and 8.2×10^7^ cells/ml, respectively, which signifies the critical cell density under high nitrogen. These values are of the same order of magnitude as the low nitrogen critical cell density (4.8×10^6^ and 5.3×10^7^ cells/ml). Between 20 and 30 hours under high nitrogen conditions we were not able to observe many cells with three bud sites (Fig. 5E) due to the cell density becoming so high that single buds were no longer visible in the images. For this reason, we did not see a statistically significant increase in the budding pattern between 0 to 10 hours and 20 to 30 hours.

Media from the time series experiment under high nitrogen conditions were also analysed by ^1^H NMR to obtain compound concentrations over time. A list of the 30 compounds identified in our high nitrogen samples and their concentrations can be seen in Table 2. Metabolites potentially involved in the budding pattern switch were again chosen from the identified compound list as those that were not present as nutrients in the raw media and had concentrations that increased during growth. Here the metabolite concentrations which corresponded to the observed switch in budding pattern were 1-propanol (1.4 – 3.4 µM), 2-hydroxyisovalerate (0.4 – 0.5 µM), 2-oxoglutarate (1.5 – 2.0 µM), 2-phenylethanol (0.4 – 0.6 µM), acetaldehyde (2.0 – 18.0 µM), acetate (27 - 186 µM), acetoin (1.0 – 5.6 µM), ethanol (472 – 4972 µM), fumarate (0.3 – 0.4 µM), isoamylol (0.7 – 0.9 µM), isopropanol (0.3 – 2.9 µM), lactate (33-231 µM), malate (2.1- 2.2 µM), pyruvate (11.5 – 38.8 µM), succinate (0.9 – 1.6 µM), and tyrosol (0.3 – 0.6 µM).

**TABLE 2:** Heatmap of the physiological concentrations of compounds detected under high nitrogen conditions during growth of *S. cerevisiae* strain Σ1278b.^a^

| Low Concentration Metabolites ($\mu\text{M}$) | | | | |
| --- | --- | --- | --- | --- |
| Metabolite Name | 0 hrs | 10 hrs | 20 hrs | 30 hrs |
| 1-Propanol | $1.3 \pm 0.1$ | $1.4 \pm 0.1$ | $3.4 \pm 0.4$ | $21.4 \pm 10.5$ |
| 2-Hydroxyisovalerate | $0.3 \pm 0.1$ | $0.4 \pm 0.1$ | $0.5 \pm 0.2$ | $1.8 \pm 0.3$ |
| 2-Oxoglutarate | $0.7 \pm 0.2$ | $1.5 \pm 0.1$ | $2.0 \pm 0.1$ | $13.6 \pm 3.2$ |
| 2-Phenylethanol | $0.4 \pm 0.1$ | $0.4 \pm 0.1$ | $0.6 \pm 0.1$ | $0.9 \pm 0.4$ |
| 3-Methylxanthine | $6.9 \pm 0.3$ | $7.8 \pm 0.1$ | $8.3 \pm 0.4$ | $6.7 \pm 1.1$ |
| Acetaldehyde | $1.5 \pm 0.4$ | $2.0 \pm 0.6$ | $18.0 \pm 5.3$ | $18.8 \pm 8.8$ |
| Acetoin | $1.2 \pm 0.1$ | $1.0 \pm 0.1$ | $5.6 \pm 1.9$ | $25.3 \pm 15.4$ |
| Choline | $2.2 \pm 0.6$ | $3.9 \pm 1.0$ | $3.7 \pm 1.0$ | $2.1 \pm 0.8$ |
| Diacetyl | $4.5 \pm 0.1$ | $4.5 \pm 0.1$ | $4.5 \pm 0.2$ | $4.2 \pm 0.1$ |
| Fumarate | $0.2 \pm 0.1$ | $0.4 \pm 0.1$ | $0.3 \pm 0.1$ | $4.8 \pm 0.6$ |
| Hydroxyacetone | $8.1 \pm 0.1$ | $7.9 \pm 0.2$ | $8.1 \pm 0.3$ | $8.4 \pm 0.2$ |
| Isoamylol | $0.8 \pm 0.1$ | $0.7 \pm 0.1$ | $0.9 \pm 0.1$ | $13.0 \pm 6.6$ |
| Isopropanol | $0.3 \pm 0.1$ | $0.3 \pm 0.1$ | $2.9 \pm 0.8$ | $2.5 \pm 0.8$ |
| Malate | $3.0 \pm 0.3$ | $2.1 \pm 0.5$ | $2.2 \pm 0.2$ | $6.2 \pm 1.1$ |
| Methionine | $0.2 \pm 0.1$ | $0.3 \pm 0.1$ | $0.5 \pm 0.1$ | $6.8 \pm 3.8$ |
| Myoinositol | $51.3 \pm 0.9$ | $50.3 \pm 0.7$ | $48.2 \pm 1.6$ | $43.7 \pm 4.2$ |
| Niacin | $9.4 \pm 0.3$ | $9.4 \pm 0.3$ | $11.6 \pm 0.9$ | $6.5 \pm 1.7$ |
| Pantothenate | $4.9 \pm 0.1$ | $5.3 \pm 0.3$ | $5.3 \pm 0.2$ | $5.5 \pm 0.2$ |
| Pyridoxine | $8.9 \pm 0.2$ | $9.0 \pm 0.3$ | $9.6 \pm 0.2$ | $10.5 \pm 1.1$ |
| Pyruvate | $8.5 \pm 0.8$ | $11.5 \pm 0.5$ | $38.8 \pm 6.3$ | $92.2 \pm 48.2$ |
| Succinate | $1.4 \pm 0.2$ | $0.9 \pm 0.1$ | $1.6 \pm 0.1$ | $18.5 \pm 10.9$ |
| Trehalose | $95.3 \pm 1.5$ | $94.4 \pm 2.3$ | $92.0 \pm 1.1$ | $64.1 \pm 18.6$ |
| Tyrosol | $0.4 \pm 0.1$ | $0.6 \pm 0.1$ | $0.3 \pm 0.1$ | $0.4 \pm 0.1$ |
| Xanthine | $1.1 \pm 0.2$ | $1.3 \pm 0.1$ | $1.2 \pm 0.4$ | $0.9 \pm 0.2$ |

| High Concentration Metabolites ($\mu\text{M}$) | | | | |
| --- | --- | --- | --- | --- |
| Metabolite Name | 0 hrs | 10 hrs | 20 hrs | 30 hrs |
| Acetate | $9 \pm 1$ | $17 \pm 1$ | $160 \pm 6$ | $411 \pm 12$ |
| Ethanol | $189 \pm 40$ | $369 \pm 8$ | $5403 \pm 160$ | $14334 \pm 1023$ |
| Formate | $221 \pm 2$ | $238 \pm 2$ | $231 \pm 9$ | $193 \pm 4$ |
| Glucose ($\times 10^3$) | $107 \pm 1$ | $103 \pm 1$ | $115 \pm 5$ | $99 \pm 1$ |
| Lactate | $8 \pm 2$ | $29 \pm 1$ | $224 \pm 8$ | $343 \pm 38$ |

<sup>a</sup>Metabolites identified through $^1\text{H}$ -NMR and quantified using the SigMa programme (Khakimov et al. 2020). Values are expressed as mean ± SE for two or three biological replicates.

Comparing these values to the concentration of metabolites produced under low nitrogen conditions at 20 hours of growth, the alcohols 2-phenylethanol and tyrosol were 60% and 63% lower under higher nitrogen respectively, while 1-propanol, isoamylol, and isopropanol concentrations were equivalent. The concentrations of some acids were also lower under high nitrogen conditions, 2-hydroxyisovalerate (↓62%), 2- oxogluatare (↓79%), fumarate (↓73%), malate (↓68%) and succinate (↓87%). However, acetate and pyruvate were 16% and 27% higher under high nitrogen conditions, respectively. Thus, while the same budding pattern behaviour was observed under different nitrogen conditions, there were significant differences in the metabolome. Collectively, these findings concur with our findings mentioned above that physiological metabolite concentrations are not responsible for the observed increase in unipolar budding.

### Proposed signalling molecule, 2-phenylethanol, can induce unipolar budding at high concentrations

Exogenous 2-phenylethanol has been reported to induce cellular signalling in *S. cerevisiae* (18, 21), but at much higher concentrations than those observed in growth media. To investigate the effects of these non-physiological concentrations, we added to low nitrogen media a concentration of 2-phenylethanol 13 times higher than those produced by the cells in our growth experiments. This higher concentration was used in previous studies to stimulate filamentous growth (18). Here we observed 91% unipolar budding for two bud sites (Fig. 4A) and 43% for three bud sites (Fig. 4B). These proportions of unipolar budding corresponded to the higher percentage of unipolar budding observed in both time series experiments between 10 to 20 hours; 84-91% for two bud sites (Fig. 3D and 5D) and 37-43% for three bud sites (Fig. 3E and 5E). These results indicate that a high non-physiological concentration of 2-phenylethanol can recapitulate the increase in unipolar budding, while physiological concentrations do not.

## DISCUSSION

The transition to filamentous growth has been proposed as a cell density dependent, chemically mediated switch in *S. cerevisiae*. This conclusion was drawn from studies which examined the growth of yeasts, metabolites, and exogenous compounds, but work has not conclusively shown the dependency of cellular density, induction, and accumulation of a chemical signal (25).

The objective of this study was to address the research gaps in the transition to filamentous growth in *S. cerevisiae.* Specifically, we focused on the switch to filamentous growth at a critical quorum sensing molecule concentration corresponding to a critical cell density, and that the intercellular signalling molecule concentrations used to induce the response were physiologically relevant (25). To achieve this, we designed a new methodology to correlate the filamentous growth of *S. cerevisiae* to both cell density and metabolite concentration. We combined chemical cell immobilisation in liquid media, automated time-lapse scanning microscopy, cell counting and ^1^H NMR. Filamentous yeast cells show a significantly higher proportion of unipolar budding overall compared to non-filamentous cells (2, 5). Thus, by using changes in budding pattern as a proxy for filamentous growth, this study was able to identify the critical cell density at which the cells switched morphology and to correlate this to the physiological concentrations of metabolites present.

Regardless of the nitrogen condition, both *S. cerevisiae* strains were in their exponential growth phase during the first 20 hours of the experiments after which their growth rate slightly declined (Fig. 3A and 5A). This trend was more pronounced for the Σ1278b strain under nitrogen deficiency (Fig. 3A). This general decline in growth is likely due to the accumulation of toxic metabolites that occurred toward the end of the experiment and nutrient starvation.

The use of the oCelloScope^TM^ for imaging enabled us to clearly observe the budding pattern of single yeast cells over time. Previous work has applied this imaging technique to bacteriological studies on real-time detection of bacterial growth and antimicrobial susceptibility (34), and to quantify filamentous bacteria (35). The methodology of monitoring yeast budding patterns presented in this paper is unique since previous studies have used photomicrographs of developing pseudohyphae in colonies growing on agar (2) or calcofluor staining of bud scars (5, 12). However, growth on agar plates does not allow the monitoring of metabolite production, budding pattern, and cell density simultaneously over time (2, 9, 18), and calcofluor staining does not permit reliable assignment of the birth end of cells (5, 11).

The highest proportion of unipolar budding measured here for cells with two bud sites was 85-91% (Fig. 3D and 5D). This finding is in accordance with previous studies that observed >85% unipolar buds for filamentous cells (2, 5). Conversely, in cells with three bud sites we found a maximum of 36-43% unipolar budding (Fig. 3E and 5E). Even though we saw a significant increase over time, this proportion was relatively low compared to the expected 97% from previous findings (5). A possible explanation for this might be that prior studies have not always reported on the position of third bud sites (2) and/or have tended to concentrate on cells at the periphery of the colony (2, 5). Our experiment analysed the whole population of cells which could explain this lower proportion. Additionally, our cells were in the exponential phase of growth and growing in liquid culture, while previous studies used cells in the stationary phase (18) or investigated the budding pattern of colonies growing on agar (2, 5).

*S. cerevisiae* strain S288c has a mutation in a gene required for filamentous growth (FLO8)(36) and therefore acted as our negative control for filamentation. Additionally, it is a haploid and thus expected to bud in a consistently axial manner (12, 13). Our findings are consistent with this since we observed S288c to bud axially at all time points under both high and low nitrogen conditions. Additionally, as anticipated for a diploid able to display filamentous growth, we observed strain Σ1278b to bud only in a bipolar or unipolar way. Furthermore, Σ1278b was observed to increase in unipolar budding, which suggests a transition to filamentous growth (2, 12–15). Our study is the first to identify the cell density at which the cells switch to a filamentous budding pattern. We define this cell density as the critical cell density, and when converted to local cell densities, we find it to be in the range of 4.8×10^6^ and 5.3×10^7^ cells/ml under low nitrogen conditions. Local cell densities are a more accurate representation of the cell density experienced by the cells in our experiments, as all cells were immobilised to the bottom of the wells.

We sought to obtain the physiological metabolite concentrations at which the cells switch to filamentous growth. For our growth phases, these were the metabolite concentrations present between 10 and 20 hours of incubation. We were able to correlate this physiological concentration of metabolites present with the critical cell density and identified a total of 29 compounds in our media (Table 1). This is comparable to other metabolome yeast studies using ^1^H NMR which identified 18-39 total compounds (37–41). The actual number of metabolites produced by *S. cerevisiae* is likely to be in the range of 600-1000 (42, 43). Our observed metabolome may be somewhat limited by the sensitivity of NMR spectroscopy. It should be noted, however, that the highest number of identified compounds achieved using currently available techniques, such as GC-TOF-MS, ranges from 80 (44) to 110 (45), so much more research needs to be done before having any overview of the full yeast metabolome and the physiological concentrations present when cells switch to filamentous growth.

Experiments to probe the mechanisms of the observed change in budding pattern were carried out with the Σ1278b strain under low nitrogen conditions. These conditions were chosen for comparison with previous studies that have used nitrogen deficiency to induce filamentous growth (2, 9, 18). Here, to investigate the effects of cell density and metabolites independently, an overnight culture growing in YPD broth was directly diluted to an inoculation concentration that corresponded to the critical cell density (Fig. 7A). In contrast to the time series experiment (Fig. 6A), there was no previous cell growth and metabolization for 10 hours in the media. Collectively, results from these experiments suggest that neither the critical cell density alone, nor a combination of critical cell density and physiological metabolites, could recapitulate the increase in unipolar budding pattern (Fig. 4A and B). The difference in the growth environment and metabolism between the different experiments could explain why we saw no change in budding pattern in either of these conditions. The observed change in budding pattern seems therefore to be linked to the specific growth environment found in the time series experiment. Additionally, when comparing metabolite concentrations under nitrogen rich and deficient conditions we see there are significant differences (Table 1 and 2). This provides extra support to the conclusion that the measured metabolites are not triggering the increase in unipolar budding.

**FIG 6.**
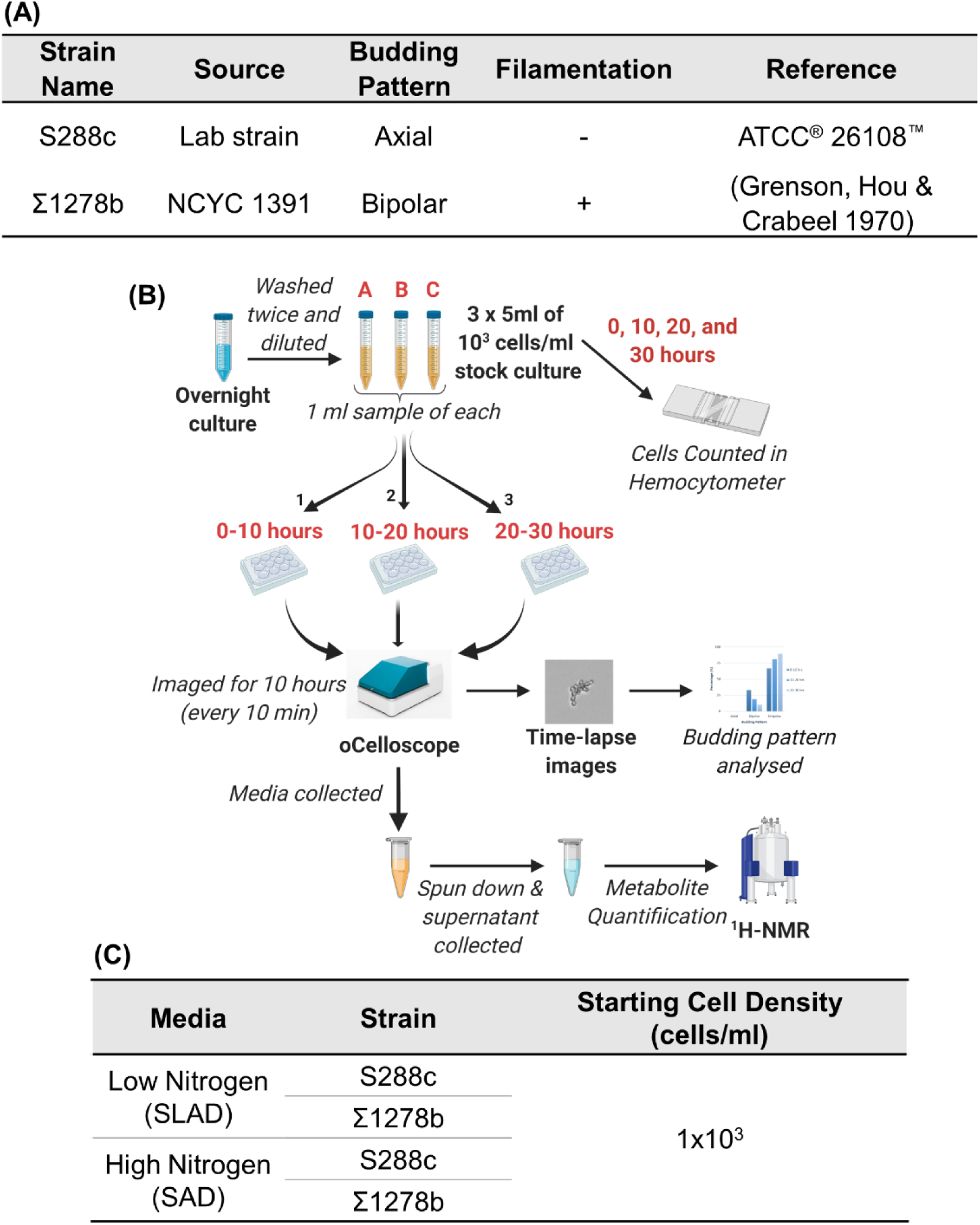
The budding patterns of *S. cerevisiae* cells growing in low and high nitrogen liquid media were tracked during growth and correlated to cell density and metabolite concentration. (A) *S. cerevisiae* strains used in this study. (B) Schematic of the time series experiment that utilised the oCelloScope^TM^ for image acquisition and ^1^H NMR for metabolite identification and quantification. (C) Experimental conditions used in the time series experiment.

**FIG 7.**
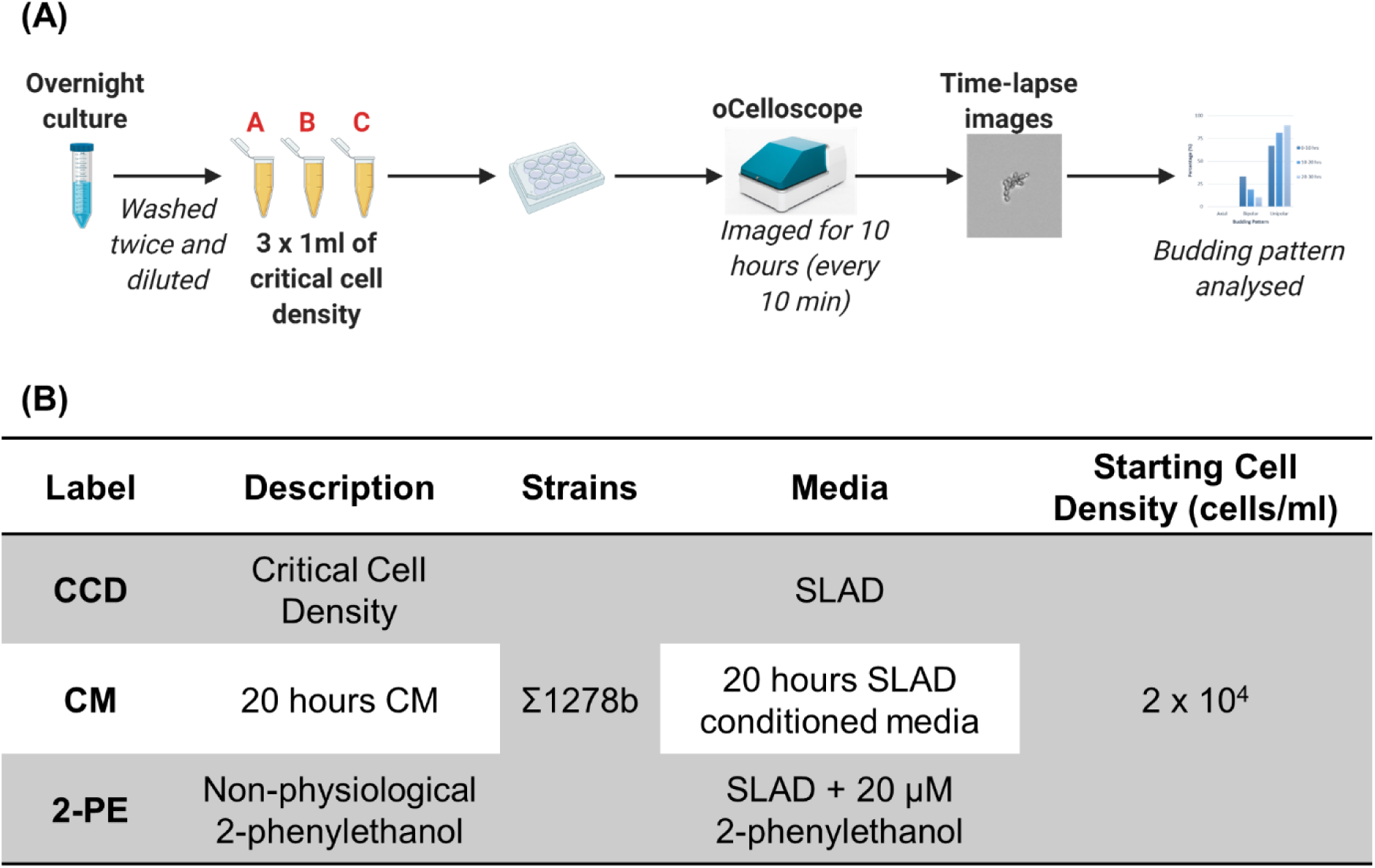
The factors responsible for change in budding pattern were investigated under low nitrogen conditions using strain Σ1278b. (A) Schematic diagram of experimental set-up. (B) Experimental conditions used included the critical cell density (CCD), 20 hours conditioned media (CM), and media with a high concentration of 2-phenylethanol (2PE). All conditions inoculated cells at the critical cell density 2 x 10^4^ cells/ml.

Some small molecule alcohols have been associated with induction of filamentous growth through a quorum sensing mechanism whereby their production is autoinduced under low nitrogen. The identified alcohols are tyrosol, 2-phenylethanol and tryptophol (18, 21–23). We were unable to demonstrate a correlation between these alcohols and the change in budding pattern. Both tyrosol and 2-phenylethanol did not show a significant increase in concentration until 30 hours under low nitrogen conditions and had very low concentrations prior to that (Table 1). Since the increase in unipolar budding occurred between 10 and 20 hours (Fig. 3D and E) we concluded that tyrosol and 2-phenylethanol could not induce the change in budding pattern. Furthermore, tryptophol was not identified in our samples, though we specifically looked for it during the NMR metabolite identification stage. Tryptophol could have been present but at very low concentrations and non detectable to the ^1^H NMR. Avbelj et al. (23) found that tryptophol was only detectable at a conc of less than 5 µM when cells reached approx. 1.5×10^7^ cells/ml. Since our cells ended their growth at a lower cell density than this, it is likely that tryptophol was present but at a non-detectable concentration. Finally, all three alcohols have relatively low volatility (2-phenylethanol bp_760_ = 220°C (46), tyrosol bp_760_ = 325°C (47) and tryptophol bp_760_ = 360°C (48)). Thus, the measured concentrations were not likely to be inaccurate due to loss during the experiment. The accumulated evidence in this paper indicates that these alcohols are not responsible for triggering the increase in unipolar budding. The discrepancy with previous studies may be attributed to differences in experimental design and execution. For example, our experiment was carried out in liquid media for 30 hours instead of colonies growing on agar plates for 3 - 5 days (9, 18, 21).

Nitrogen deficiency has commonly been linked to inducing filamentous growth in *S. cerevisiae* (2, 18). Contrary to expectations, under both nitrogen rich and deficient conditions we see a significant increase in unipolar budding between 10 to 20 hours of growth (Fig. 3B-E, 5B-E) occurring at local cell densities of a similar order of magnitude (between 5.2×10^6^ and 8.2×10^7^ cells/ml under high nitrogen and 4.8×10^6^ and 5.3×10^7^ cells/ml under low nitrogen). Therefore, nutrient deficiency alone appears not to trigger the switch to unipolar budding and perhaps the switch relies on other factors.

The results presented in this study have important implications in terms of satisfying the necessary criteria to be classified as a quorum sensing mechanism; specifically the criteria that filamentous growth is triggered at a critical cell density and is reproducible at physiological signal molecule concentrations (25). Therefore, our findings that the critical cell density and physiological metabolite concentration are not responsible for the shift to a filamentous budding pattern leads to the conclusion that this phenomenon is not occurring through a quorum sensing mechanism but is being controlled in an alternative way. While the change in budding pattern observed in the time series experiment appears to be cell density dependent, it does not however align with an authentic quorum sensing mechanism based on the previously defined criteria (25). It is only when the cells could increase in cell density in the same media over time that this shift to unipolar budding was observed. These findings raise intriguing questions regarding the mechanism of action controlling this transition to filamentous growth and highlight the need for further studies into this important issue.

An alternative mechanism could be one involving cell-to-cell-contact that was found to be a mechanism of growth arrest in yeast (49, 50). Cellular communication mediated by direct cell-to-cell contact has also been found in bacteria. For example, direct cell-to-cell contact results in growth inhibition in *Escherichia coli* (51), coordinates cellular motility in Mycobacterium (52), and mediates auto aggregation and adhesiveness in *Lactobacillus acidophilus* cell surface (53). Thus, there may be a signal present on the cell surface, only present there because of metabolism, that signals the change in budding pattern. Further investigation is therefore required to determine the mechanism by which the cells increase in unipolar budding since there appears to be more transpiring than a quorum sensing mechanism.

As mentioned above, 2-phenylethanol has previously been linked to inducing filamentous growth under low nitrogen conditions (18, 21). In these previous experiments, the compound was added at concentrations much higher than those produced by the cells growing in defined media (18, 21, 22). In our experiments, a high non-physiological concentration of 2-phenylethanol, i.e. 20 µM compared to the maximum of 4.3 µM measured in our low nitrogen media, induced an increase in unipolar budding whereas physiological levels of both 2-phenylethanol and other metabolites did not. This finding confirms the association between 2-phenylethanol and filamentous growth. However, an implication of the non-physiological concentration used is that 2-phenylethanol-induced filamentous growth cannot be occurring through a quorum sensing mechanism as previously suggested (18, 23). Table 3 compares our 2- phenylethanol findings with the criteria defined by Winters et al. (25) and finds that it does not satisfy two key points. Therefore, we hypothesise that 2-phenyethanol induces filamentous growth through a toxicity mechanism due to its high concentration rather than through an intercellular signalling mechanism.

**TABLE 3.** 2-phenylethanol does not act as an intercellular signaling and quorum sensing molecule to induce the change in budding pattern based on our experimental results.^a^

| *Criteria* |  | **2-phenylethanol** |
| --- | --- | --- |
| ***Intercellular Signalling Molecule*** | Extracellular & Identified | Y |
|  | Specific signalling mechanism | Y |
|  | Reproducible response | N<sup>b</sup> |
|  | Adaptive at community level | Y |
| ***Quorum Sensing Molecule*** | Proportional to cell density | Y |
|  | Critical threshold trigger | N<sup>c</sup> |

<sup>a</sup> Criteria from (23).
<sup>b</sup> Response only observed when 2-phenylethanol used at much higher concentrations (20 µM) than found to be physiologically produced (0.7 – 1.5 µM)
<sup>c</sup> Since the response is only induced at high levels it may be a starvation or stress response and not a QS mechanism

A recent study by Lenhart et al. (54) also provides evidence contrary to previous assumptions of quorum sensing mechanisms in *S. cerevisiae.* They found that the induction of morphogenesis by high concentrations of 2-phenylethanol and tryptophol (100 µM) is not observed in most environmental isolates and that the filamentous induction response in Σ1278b was small compared to the extent of that reported by Chen and Fink (18). This supports our conclusion that quorum sensing controlled filamentous growth in yeast is not straightforward and that the mechanism of this biologically critical phenomenon requires more examination.

In conclusion, the methodology presented in this paper uses budding pattern as a proxy for filamentous growth. This study monitors the budding pattern of yeast cells over time while simultaneously collecting data on metabolite concentration and cell density. This allowed us to probe the question of whether filamentous growth is induced though a quorum sensing mechanism based on more specific criteria. We identified the critical cell density and physiological metabolite concentration present during the shift to a filamentous budding pattern in *S. cerevisiae*. However, neither cell density, metabolite concentration nor nitrogen condition triggered the increase in unipolar budding. An implication of these findings is that the criteria for filamentous growth to occur via an intercellular signalling and quorum sensing mechanism were not fulfilled and that the change in budding pattern does not therefore occur through these mechanisms. Our results also indicated that 2-phenylethanol induces filamentous growth through another mechanism, proposed as toxicity, and not through a quorum sensing mechanism as previously accepted.

The contribution of this study has been to provide clarity of this transition in the model and industrially useful species, *S. cerevisiae*, highlighting the need for further investigation to determine the mechanism controlling the shift to filamentous growth under physiological conditions. It thus challenges the assumptions around quorum sensing in *S. cerevisiae* widely accepted in literature. The mechanism by which the switch to filamentous growth occurs in *S. cerevisiae* needs to be elucidated through further research without resorting to categorisation into quorum sensing mechanisms.

## MATERIALS AND METHODS

### Yeast Strains and Media

The *Saccharomyces cerevisiae* strains S288c and Σ1278b were used as a negative and positive control for filamentation respectively (Fig. 6A). The strains were maintained using accepted protocols, detailed below.

Yeast Extract Peptone Dextrose (YPD) broth was made by combining 1 g yeast extract (BD Bacto^TM^, Herlev, Denmark), 2 g bacteriological peptone and 2 g glucose per 100 ml of distilled water. The solution was stirred with heating until dissolved. 2 % YPD agar plates were made in the same way but with addition of 2g/100ml bacteriological agar to the YPD broth before autoclaving. Synthetic Low Ammonium Dextrose (SLAD) broth was made by combining 0.67 g Yeast Nitrogen Base without amino acids and ammonium sulphate (BD Difco^TM^, Herlev, Denmark), 2 g glucose and 50 µl of ammonium sulphate solution (1M) per 100 ml of distilled water. Synthetic Ammonium Dextrose (SAD) liquid broth was made by combining 0.67 g Yeast Nitrogen Base without amino acids and ammonium sulphate, 2 g glucose and 3.7 ml of ammonium sulphate solution (1M) per 100 ml of distilled water. All media were autoclaved and cooled to room temperature and sterile filtered (Q-Max 25 mm 0.22 µm CA) before use. Concanavalin A (biotin conjugate, Type IV) at 1mg/ml MilliQ water (Con A) was used for chemical immobilization of cells. Where not indicated otherwise, all chemicals were purchased from Sigma–Aldrich (Søborg, Denmark).

### Image Acquisition

Images of growing cells in liquid media were obtained using an oCelloScope^TM^ (BioSense Solutions ApS, Farum, Denmark). The oCelloScope^TM^ is a digital time-lapse microscopy technology that generates very clear images of growing microorganisms. This is achieved by scanning through a sample to produce a series of images using a tilted imaging plane. Combining the tilted images give the best focus images and enables time-lapse images of single cells (34, 35).

### Time series determination of budding pattern under low and high nitrogen conditions

S288c and Σ1278b strains of *S. cerevisiae* (Fig. 6A) were used in the time series experiment which linked budding pattern with cell density and physiological metabolite concentrations. Figure 6B represents a schematic of the overall time series experimental process used in this research. The experiment was carried out in both high (SAD) and low nitrogen (SLAD) liquid media with the same starting cell density of 10^3^ cells/ml (Fig. 6C). The overnight culture was made by inoculating 5 ml of YPD broth in a 15 ml centrifuge tube with a single yeast colony grown on a 2% YPD agar plate. This was incubated overnight at 25°C with shaking. After overnight incubation, the cells were counted using a hemocytometer (Neubauer Improved counting chamber, Assistent), washed twice with the relevant media, and diluted to obtain a cell density of 1×10^3^ cells/ml in the relevant media. Three 5 ml stock cultures were made in 15 ml centrifuge tubes with lids to produce biological replicates A, B and C for each condition (Fig. 6C). During the experiment these stock cultures grew at 25°C without shaking. Wells of the 12-well microtiter plate (VWR 734-2324 T C-treated 12-well plates) were prepared just before inoculation and imaging by coating the bottom of each well with 15 µl of Con A and allowing the wells to dry. 1 ml of each stock culture was pipetted into wells of the microtiter plate. After inoculation with the cell suspension the plate was placed in the oCelloScope^TM^ and allowed to rest for 30 minutes before starting image acquisition. This was to ensure that the cells were all immobilized on the bottom of the well before imaging to aid focusing. Scanning was set to take place every 10 minutes for 10 hours with one scan area per well at 25°C. Two scan areas per well were used for the final 20-to-30-hour time point to ensure enough single cells were captured at the higher cell density. After 10 hours of image acquisition, the plate was removed, and the media from each well was collected in 1.5 ml Eppendorf tubes. The media was spun down in a centrifuge at 4°C for 10 min at 4400 g (Centrifuge 5920 R, Eppendorf Nordic, Hørsholm, Denmark) and the supernatant collected and stored at -80° for ^1^H NMR analysis. Direct transfer of 1 ml of each stock solution occurred again at 10 and 20 hours to a freshly Con A coated microtiter plate, as previously described. Images at 10 and 20 hours were acquired as previously described. Thus, this process resulted in a total of 30 hours of imaging to give three image acquisitions time periods of 10 hours each. Cell density of the stock cultures was also measured at 10, 20 and 30 hours with a hemocytometer. Cell densities were averages of the three biological replicates A, B and C.

### Investigating the factors responsible for change in budding pattern

Figure 7A represents a schematic of the overall process for the experiments used to investigate the effect of various factors of interest on the budding pattern. This included investigating the effect of critical cell density, conditioned media and high 2- phenylethanol concentrations (Fig. 7B). All experiments were carried out using a cell density which corresponded to the critical cell density identified under low nitrogen conditions in the time series experiment (2×10^4^ cells/ml).

### Effect of critical cell density alone on budding pattern

To test the effect of critical cell density on the budding pattern, we analyzed the effect of using cells at the critical cell density in low nitrogen broth. The critical cell density condition used SLAD media inoculated with the critical cell density of 2×10^4^ cells/ml (Fig. 7B). 5 ml of YPD broth in a 15 ml centrifuge tube was inoculated with a single yeast colony grown on a 2% YPD agar plate and incubated overnight at 25°C with shaking. After overnight incubation, the cells were counted using a hemocytometer. These cells were then washed twice with SLAD media. A cell density of 2×10^4^ cells/ml (critical cell density) was obtained to make three 1 ml solutions (Fig. 7A). This produced three biological replicates, A, B and C. The microtiter plate was prepared as in the time series experiment. The three 1 ml solutions were pipetted into the relevant wells before analysis. Image acquisition in the oCelloScope^TM^ and image analysis to obtain the budding pattern was carried out as in the time series budding pattern assay.

### Combined effect of critical cell density and metabolites on budding pattern

To test the effect of critical cell density and metabolites on budding pattern, we inoculated the critical cell density of 2×10^4^ cells/ml into conditioned media grown for 20 hours (Fig. 7B). Conditioned media was produced by growing 10^3^ cells/ml of freshly washed cells in SLAD broth for 20 hours followed by centrifugation and supernatant removal. This conditioned media supernatant was used immediately in the experiment to wash and inoculate cells, as described in the previous section.

### Effect of non-physiological 2-phenylethanol on budding pattern

To investigate the effect of non-physiological concentrations of 2-phenylethanol on the budding pattern we used SLAD broth containing 20 µM of 2-phenylethanol. This concentration was previously found to induce filamentation by Chen and Fink (18) under low nitrogen. Cells were inoculated into the spiked broth at the critical cell density of 2×10^4^ cells/ml (Fig. 7B). Cell preparation, inoculation and imaging was carried out as described in the previous section.

### Image analysis to determine budding pattern

Manual image analysis was carried out on the images acquired from the oCelloScope^TM^ to determine the budding patterns of individual cells. Firstly, the scan areas were cropped using a custom-made code in MATLAB (version R202a, The MathWorks Inc., Natick, MA, USA) such that thirty images were cropped from each scan area from the three biological replicate wells. Samples of cropped images were randomly taken from the biological replicates A, B and C. Each cropped image was then analyzed by eye to determine the bud site sequence of new daughter cells over each period of image acquisition to obtain a bud site sequence for >200 cells and >300 bud sites for each condition (Fig. 1). This was done by observing new daughter cells and recording the positions of the first two or three bud sites (dependent on the number of times the cell budded within the time frame) to assign each cell with a bud site sequence. An example of this process on a representative cell can be found in Figure 2 (for full time-lapse see Movie S1). These were then assigned their corresponding budding pattern classification (axial, bipolar or unipolar) according to Figure 1. From the budding pattern assignments the percentage value of each budding pattern was determined for the different experimental conditions.

### Cell Density Data Analysis

As mentioned above, the cell density in the 1 ml (V_sample_) of stock solution was determined before imaging by counting with a hemocytometer. This measured cell density value (measured CD) was converted to a local cell density (local CD) that the cells would experience based on being immobilized on the bottom of the well.

Assuming a yeast cell size of 10 µm, we calculated the volume that the yeasts would occupy at the bottom of the well. Since the well had a bottom surface area of 3.85 cm^2^, the new volume occupied by the yeast at the bottom of the well would be 3.85×10^-3^ cm^3^ (V_bottom_) if the whole bottom surface was covered with yeast. This value was used to convert the cell-density per ml value to a local cell density according to the following formula:

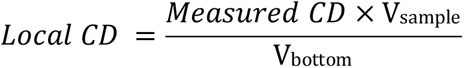

### NMR Chemicals and Buffer Preparation

Analytical grade sodium phosphate monobasic (NaH2PO3, ≥99.0%), sodium phosphate dibasic dihydrate (Na_2_HPO_3_, 2 H_2_O, ≥98.0%), sodium azide (NaN_3_, ≥99.5%), deuterium oxide (D_2_O, 99.9 atom % D), and 3-trimethyl-silyl-[2,2,3,3-2H4] propionic acid sodium salt (TSP-d4, 98 atom % D) were purchased from Sigma–Aldrich (Søborg, Denmark). Water used throughout the study was purified using a Millipore lab water system (Merck KGaA, Darmstadt, Germany) equipped with a 0.22 mm filter membrane.

### NMR Sample Preparation

An aliquot of 700 µL of sample was transferred into a 2 mL Eppendorf tube containing 300 µL of phosphate buffer (0.15 M, 20% D2O). The mixture was vigorously vortexed for 30 s. A volume of 600 μL was transferred into 5 mm O.D. NMR SampleJet tubes (Bruker Biospin, Ettlingen, Germany).

### NMR Measurements

Proton (^1^H) NMR spectra were recorded on a Bruker Avance III 600 operating at a proton’s Larmor frequency of 600.13 MHz. The spectrometer was equipped with a 5 mm broadband inverse (BBI) probe and an automated sample changer (SampleJet™, Bruker Biospin, Ettlingen, Germany). The SampleJet was equipped with a refrigerated sample storage station with cooling racks (278 K) and heating/drying station (298 K). Data acquisition and processing were carried out in the TopSpin software (version 3.5, Bruker, Rheinstetten, Germany). Automation of the overall measurement procedure was controlled by iconNMR™ (Bruker Biospin, Rheinstetten, Germany). NMR spectra were measured at 298 K using the standard pulse sequence for pre-saturation of the water signal (zgcppr pulse program, Bruker nomenclature), a 90° pulse, a sweep width of 9615.385 Hz (16 ppm), and an acquisition time of 3 s. The relaxation delay (d1) was set to 4 s. The receiver gain (RG) value was determined experimentally for each sample. NMR data was collected into 32 K data points after 512 scans. Each free induction decay was apodized by a Lorentzian line-broadening of 0.3 Hz. Automatic phase and baseline correction were performed in TopSpin.

### Processing of the NMR data and metabolites quantification

NMR spectra were imported into the SigMa software (33) where the ^1^H NMR signals were referenced to the TSP signal at 0.00 ppm. Signals alignment and quantification (relative) were performed in SigMa by icoshift and multivariate curve resolution, respectively (55, 56). The yeast metabolome database (http://www.ymdb.ca/) and the Chenomx NMR suite (Chenomx Inc. Edmonton, Canada) were used for metabolite identification.

Metabolite concentrations (absolute) were calculated using the following equation:

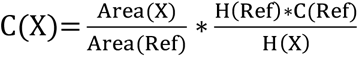

Where C(X) is the concentration of the metabolite X, Ref is the reference compound (TSP), and H indicates the number of protons giving rise to the signal from metabolite X and the reference compound (TSP). Metabolite concentrations are averages of the three biological replicates A, B and C.

Due to the short recycle delay (D1) employed for the ^1^H NMR experiments (D1=4) - too short for an accurate absolute quantification of TSP (D1≥20) - metabolite concentrations (absolute) were calculated in two-steps: firstly, the average glucose concentration in the raw media (known) was used to retrieve an estimate of the TSP concentration (absolute) in the very same samples. Secondly, the average absolute concentration of TSP was used to calculate the metabolite concentrations (absolute) in the remaining samples. Therefore, metabolites concentrations reported in the present study are to be considered as estimates of absolute concentrations.

### Statistical Analysis

Statistical analyses were performed using Minitab® 19.2020.1 (64-bit). To determine statistically significant differences between conditions of budding pattern proportions, a 2×2 contingency table using Fisher’s Exact Test at a significance level of p=0.05 was applied. For metabolite concentrations, a minimum of two biological replicates were averaged to obtain the mean, and standard deviations calculated to determine the standard error of the mean. For cell densities, the three biological replicates were averaged to obtain the mean value, and the standard deviations calculated.

## ACKNOWLEDGEMENTS

This research was supported by the Faculty of Veterinary and Agricultural Science at the University of Melbourne and an Australian Government Research Training Program (RTP) Scholarship.

We are grateful to Nadia Devargue, Chu Chu Huang and Sebastian Bech-Terkilsen from the University of Copenhagen for assistance in the laboratory, and Franciscus Winfried J van der Berg from the University of Copenhagen for allowing us use of an oCelloScope^TM^.

We declare that we have no conflict of interest.

Experiment designed by MW, KH and NA; experiment and analysis carried out by MW; NMR data collected by VA; paper written by MW. All authors have read, critically revised, and approved the final version of the manuscript.

## SUPPLEMENTAL MATERIAL

**MOVIE S1** Full representative cropped image time-lapse with bud site sequence assignment and budding pattern categorisation from Figure 2.

**TABLE S1.** Results from Nitrogen Concentration Assay which measured the nitrogen concentration from the media and the supernatant from the three biological replicates A, B and C at the end of the time series experiment i.e. after 30 hours of growth.

| Nitrogen Condition | High Nitrogen |  |  | Low Nitrogen |  |  |
| --- | --- | --- | --- | --- | --- | --- |
| Condition | Media | Σ1278b | S288c | Media | Σ1278b | S288c |
| Nitrogen Conc (mM) | 82.9 | 80.0 | 85.0 | 1.06 | 0.04 | 0.03 |

**TABLE S2.**
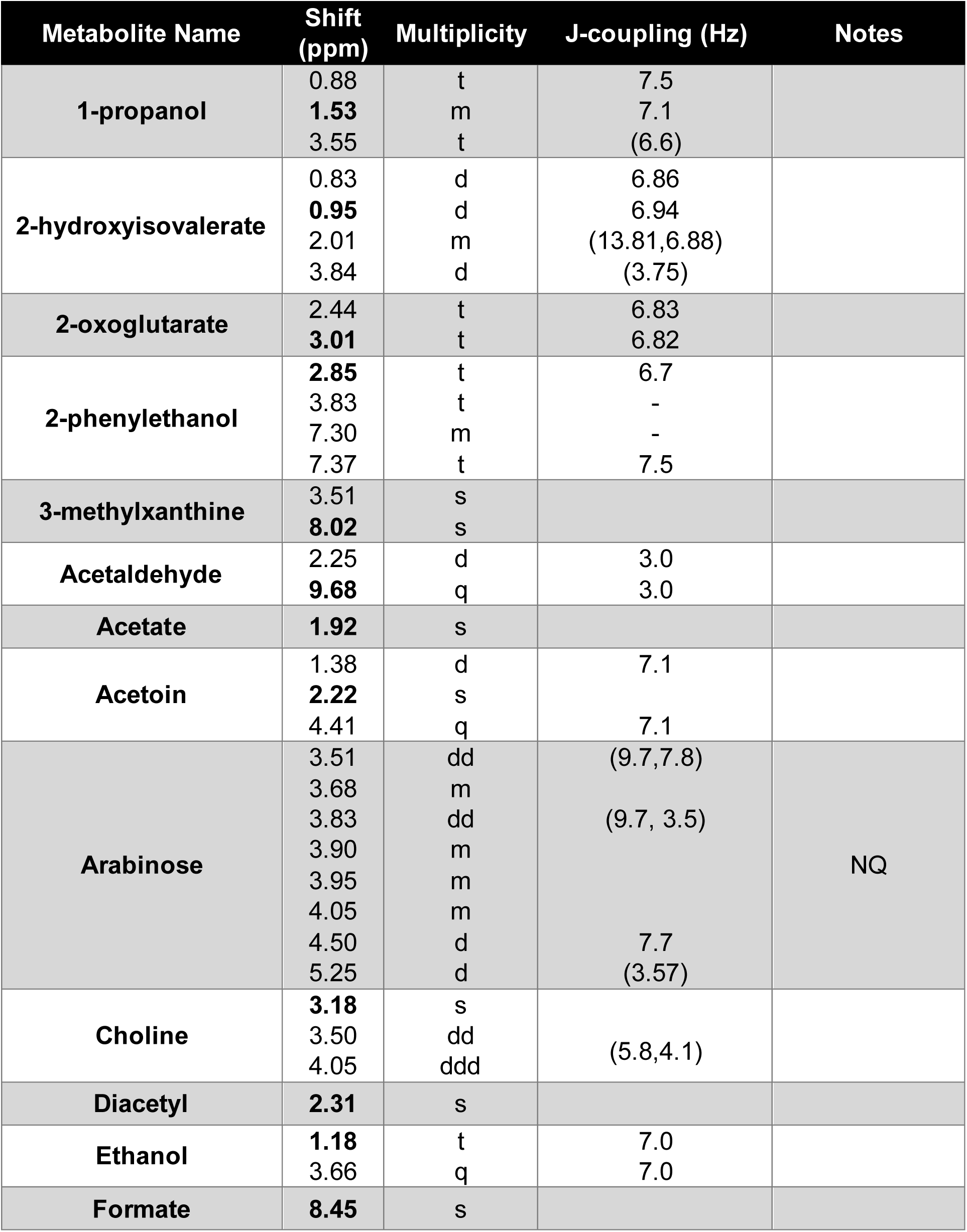
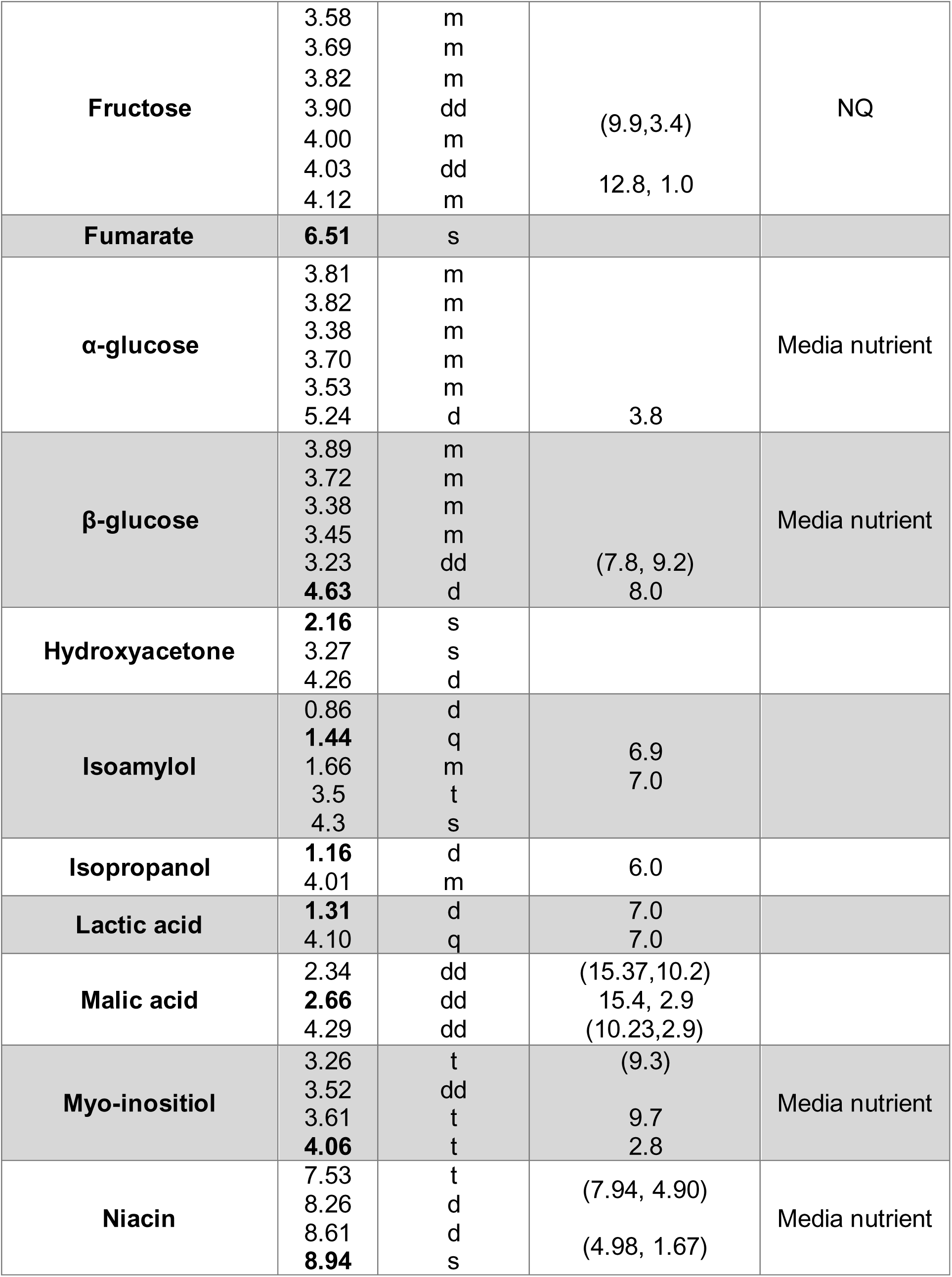
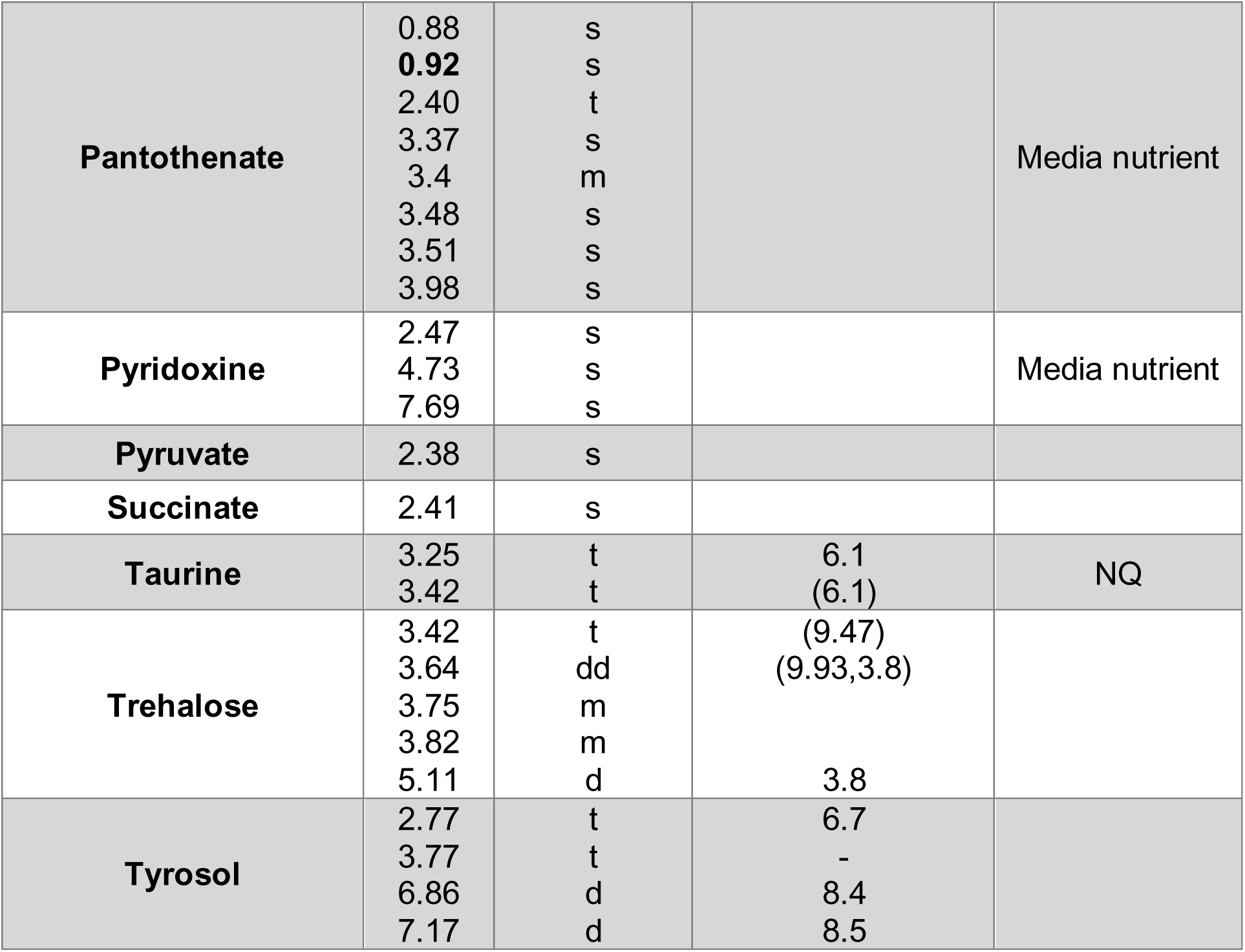
^1^H-NMR metabolite shifts and intervals with J-coupling. Shifts in bold were used for quantification.

**FIG S1.**
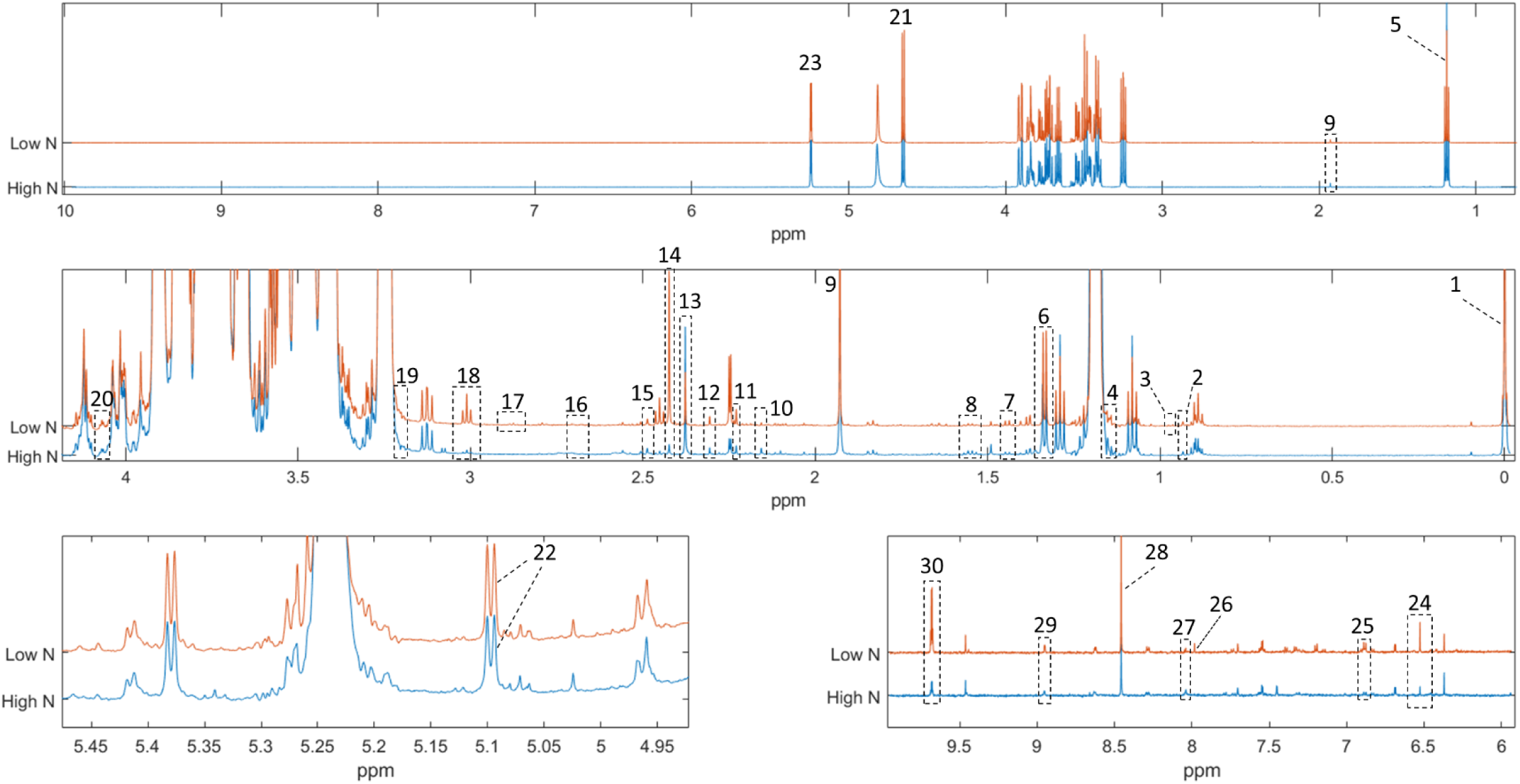
Representative ^1^H-NMR spectra of samples. The major resonances used for quantification have been assigned: 1. TSP; 2. Pantothenate; 3. 2-hydroxyisovalerate; 4. Isopropanol; 5. Ethanol; 6. Lactate; 7. Isoamylol; 8. 1-Propanol; 9. Acetate; 10. Hydroxyacetone; 11. Acetoin; 12. Diacetyl; 13. Pyruvate; 14. Succinate; 15. Pyridoxine; 16. Malate; 17. 2-Phenylethanol; 18. 2-oxoglutarate; 19. Choline; 20. Myoinositol; 21. β-Glucose; 22. Trehalose; 23. α-Glucose; 24. Fumarate; 25. Tyrosol; 26. Xanthine; 27. 3- Methylxanthine; 28. Formate; 29. Niacin; 30. Acetaldehyde.

**FIG S2.**
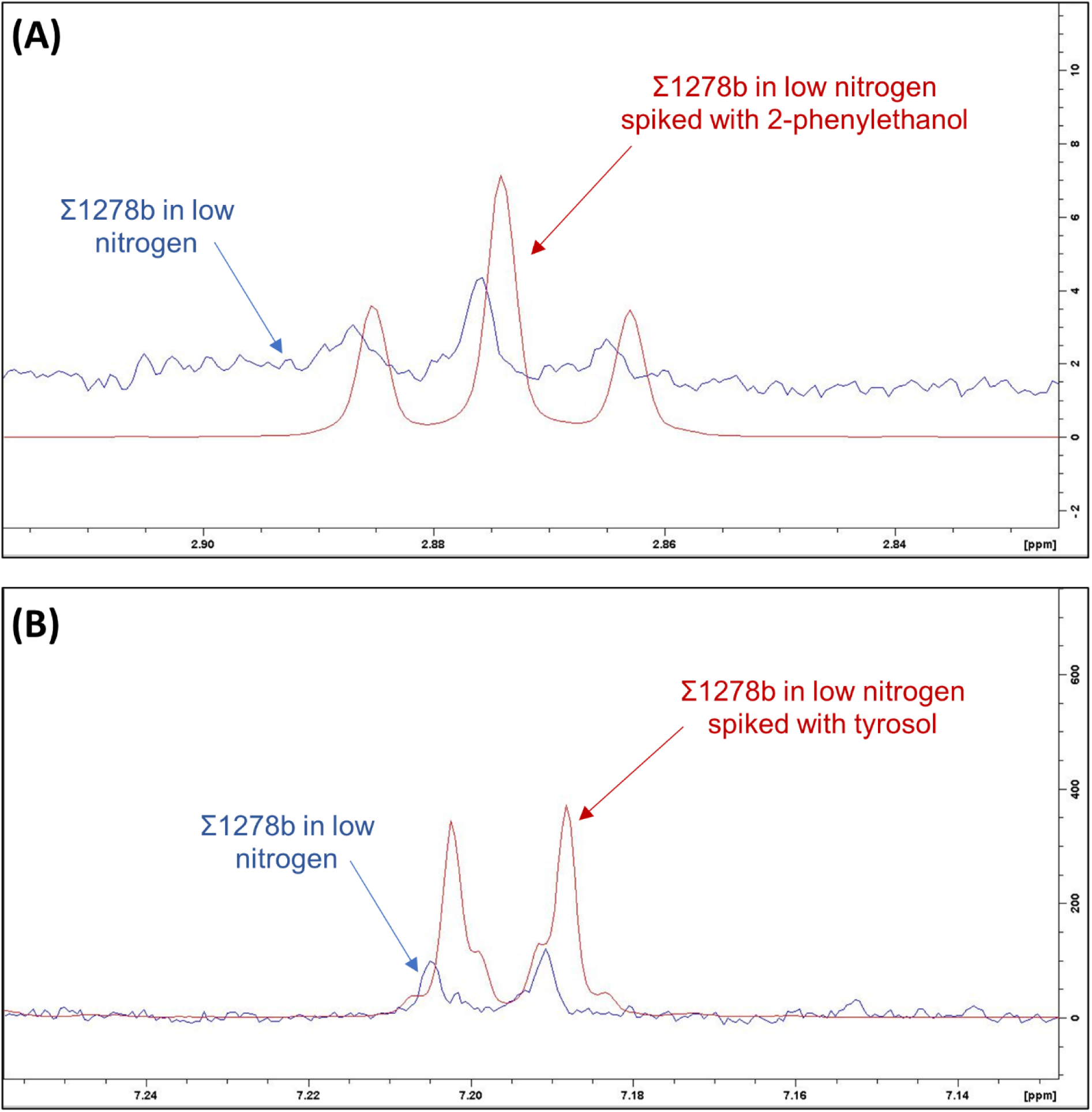
Representative ^1^H-NMR spectra showing the results of the spiking experiment conducted by adding (A) 10 µL 2-phenylethanol and (B) 5 µL tyrosol to the samples.

